# Dissolving microneedle array patches containing mesoporous silica nanoparticles of different pore sizes as a tunable sustained release platform

**DOI:** 10.1101/2023.09.27.559712

**Authors:** Juan L. Paris, Lalitkumar K. Vora, Ana M. Pérez-Moreno, Yara A. Naser, Qonita Kurnia Anjani, José Antonio Cañas, María José Torres, Cristobalina Mayorga, Ryan F. Donnelly

## Abstract

Dissolving microneedle array patches (DMAPs) enable efficient and painless delivery of therapeutic molecules across the *stratum corneum* and into the upper layers of the skin. Furthermore, this delivery strategy can be combined with the sustained release of nanoparticles to enhance the therapeutic potential in a wide variety of pathological scenarios. Among the different types of nanoparticles that can be included in microneedle formulations, mesoporous silica nanoparticles (MSNs) of tuneable pore sizes constitute a promising tool as drug delivery systems for cargos of a wide range of molecular weights. However, the development of efficient methods to produce DMAP containing large amounts of MSNs of different pore sizes has not been reported. In this work, DMAP containing MSNs with varying pore sizes was prepared and characterized. After synthesizing and characterizing MSNs, the pore size of the nanoparticles (in the range of 3 to 13 nm for S-MSN and XL-MSN, respectively) was observed to influence the loading and release of both small and large molecules, using fluorescein and ovalbumin (OVA) as model cargos. Moreover, a new preparation method was developed to produce DMAP containing large amounts of these MSNs located mainly in the microneedle tips. The successful insertion of these DMAPs was confirmed *in vitro* (using Parafilm), *ex vivo* (using excised neonatal porcine skin) and *in vivo* (in the back of mice) models. The dissolution of the microneedles and deposition of the nanoparticles inside the skin were also confirmed both *ex vivo* and *in vivo* using fluorescent nanoparticles, with complete microneedle dissolution after 2 h of insertion *in vivo*. Through histological studies, the microneedle-delivered MSNs were found to end up inside antigen presenting cells in the skin tissue (either F4/80^+^ macrophages or CD11c^+^ dendritic cells). For this reason, the uptake and biological effect of the MSNs was evaluated *in vitro* in dendritic cells, showing that while smaller pore MSNs were taken up by cells more efficiently (with over 80 % of S-MSN uptake compared to *ca*. 55 % for XL-MSNs), the dendritic cells treated with OVA-loaded XL-MSNs underwent the largest degree of activation (inducing over 25 % of CD40 expression compared to less than 2 % for OVA-loaded S- MSNs). Finally, the immune response to OVA-loaded XL-MSNs in mice was evaluated after repeated administration either subcutaneously or through DMAP. The results of this experiment showed comparable levels of anti-ovalbumin immunoglobulin generation through both routes of administration (with significant production of OVA-specific IgG1 and IgG2b antibodies), highlighting the good potential of this delivery platform for vaccination or immunotherapy applications.

## 1. Introduction

Dissolving microneedle array patches (DMAPs) are arrays of needle-like structures with microscale diameters and lengths up to 1 mm mainly made using polymers that dissolve with the interstitial fluid after their insertion into the skin[1]. Given their small size, when they are inserted in the skin, while they enable the deposition of drugs inside the upper layers of the skin (and across the external barrier of *the stratum corneum*), they do not reach blood vessels or pain receptors, not producing any bleeding or pain. For this reason, they are often proposed as an alternative drug administration option without the need for conventional injections[2–4]. DMAPs have been proposed for a wide variety of therapeutic applications, such as vaccines[5– 7], cancer treatment[8,9], migraines[10], fungal infections[11], psoriasis[12] or malaria[13], among many others. Within the polymer matrix that makes up the microneedles, different micro- or nano-particle based formulations can be included to improve the therapeutic performance of the formulation[14]. For example, if nanoparticles with sustained drug release are administered through DMAP, their deposition inside the skin creates a reservoir of the drug at the site of administration, which is slowly released from the nanoparticles, reducing the need for continuous re-administration of the treatment. Based on this concept, DMAP has been prepared containing many types of nanoparticles, such as liposomes[15–17], cubosomes[18,19], polymeric nanoparticles[20–22], metallic nanoparticles[23] and mesoporous silica nanoparticles (MSNs)[24]. Among these different types of nanoparticles, MSNs with tunable pore sizes (generally in the range of 2-20 nm in diameter[25]) constitute a particularly promising tool as drug delivery systems[25–27], as they present a large loading capacity for cargos of a wide range of molecular weights[28–30]. MSNs have been proven to be safe and to undergo dissolution in physiological environments, giving rise to nontoxic degradation products such as silicic acid, which can be safely excreted in urine[31,32]. Several microneedle arrays (either dissolving or nondissolving) containing MSNs have been previously reported[24,33–35]. However, the amount of MSNs contained in these previously reported formulations was relatively low and would likely not be enough to deliver a therapeutically relevant dose of drug for most potential applications. Furthermore, the possibility of preparing DMAP with MSNs of different pore sizes has not previously been reported to the best of our knowledge. The development of an efficient method to prepare DMAP containing large quantities of MSNs of different pore sizes would provide a tunable platform that could be adapted for many different therapeutic applications, as the pore size of the nanoparticles could be tailored for the desired drug[27], and might even allow for combination therapies in which a mixture of different MSN formulations (each optimized for a different drug) could be codelivered through a single DMAP. In this work we report for the first time a simple method to obtain DMAP containing large amounts of MSNs of tuneable mesopore sizes. Different nanoparticle-containing DMAP formulations were prepared and characterized, and their therapeutic potential was assessed through a variety of *in vitro, ex vivo* and *in vivo* methods. The modular platform presented here could be adapted to deliver sustained release formulations of therapeutic molecules over a wide range of molecular weights, either as monotherapies or as combination therapies with multiple drugs.

## 2. Materials and Methods

### 2.1 Materials

The following reagents were purchased from Merck (Sigma−Aldrich, Spain) and were used without further purification: tetraethylorthosilicate (TEOS), cyclohexane, triethanolamine, cetyltrimethylammonium chloride (CTAC), ammonium nitrate, ethanol, hydrochloric acid, fluorescein isothiocyanate (FITC), rhodamine B isothiocyanate (RITC), aminopropyltriethoxysilane (APTES), fluorescein sodium salt, ovalbumin (OVA), phosphate buffered saline (PBS) tablets, polyvinyl alcohol 9-10 kDa (PVA) and polyvinyl pyrrolidone K 30 (PVP, 40 kDa), agarose, Roswell Park Memorial Institute (RPMI)-1640 culture medium, fetal bovine serum, nonessential amino acids, L-glutamine, and β-mercaptoethanol. Antibodies and other reagents for immunofluorescence and ELISA experiments were obtained from Thermo Fisher Scientific (Spain). FITC-labelled anti-mouse CD40 for flow cytometry was purchased from Biolegend (Spain).

### 2.2 Synthesis of MSNs

MSNs of different pore sizes were prepared by a previously described biphasic method based on the condensation of TEOS in a biphasic water/cyclohexane system, using triethanolamine as the base and CTAC as the structure-directing agent surfactant[25,26]. The aqueous phase was composed of a mixture of 24 mL of a commercial aqueous solution of CTAC (25% w/v)), 0.18 g of triethanolamine and 36 mL of deionized water. The organic phase consisted of 20 mL of a mixture of cyclohexane with TEOS. The concentration of TEOS depended on the material to be prepared: 40% for S-MSNs, 20% for M-MSNs, 10% for L-MSNs and 5% for XL-MSNs. The synthesis reaction was carried out at 50°C for 24 h. Then, the surfactant was extracted by ion exchange with an ethanolic solution of ammonium nitrate (10 mg/mL) at reflux for 1 h, followed by a second reflux for 2 h in an ethanolic solution of 12 mM HCl. Finally, the material was washed with ethanol 3 times to obtain the desired materials, which were dried and stored at room temperature until further use. Fluorescent MSNs were also obtained by adding a mixture of 1.5 mg of fluorescein FITC or RITC and 15 μL of APTES in 1 mL of ethanol in the aqueous phase during MSN synthesis.

### 2.3 Cargo loading and release from MSNs

Fluorescein sodium salt (as a model for small molecule drugs) or OVA (as a model for therapeutic proteins) was loaded in MSNs by dispersing 10 mg of MSNs in a 10 mg/mL solution of the cargo in PBS (10 mM, pH=7.4) and stirring overnight. Then, the loaded particles were collected by centrifugation, and the nonloaded cargo was quantified from the supernatant by UV−Vis spectrophotometry. For release experiments, loaded particles were suspended in PBS and stirred at 37°C. At different time points, the particles were centrifuged, released cargo was quantified by fluorimetry (fluorescein sodium salt, λ_EX_=580 nm; λ_EM_=520 nm) or UV−Vis spectrophotometry (OVA, λ_ABS_=280 nm), and the particles were resuspended in fresh PBS to continue stirring at 37°C.

### 2.4 Characterization techniques

Dynamic light scattering (DLS) and Z-potential measurements were performed with a Malvern Zetasizer Nano ZS90 instrument, checking both particle size and surface charge. The instrument used was equipped with a “red laser” (λ = 300 nm), and DLS measurements were performed with a detection angle of 90°, while the Smoluchowski approximation was used for Z-potential measurements. To check the morphology and the different pore sizes of the nanoparticles, the characterization of the nanoparticles was performed by transmission electron microscopy (TEM) on a Thermo Fisher Scientific Tecnai G2 20 Twin using copper grids of mesh size 200 coated with a Formvar-Carbon film. Scanning electron microscopy (SEM) was carried out on an FEI Quanta- 250 microscope (Thermo Fisher Scientific, USA) after coating the samples with a thin layer of gold under vacuum. Nitrogen adsorption (in a Micromeritics ASAP 2020 unit) measurements were carried out at the Central Research Support Services (SCAI) of the University of Malaga (UMA). Fluorimetry and UV−Vis spectrophotometry were carried out using a plate reader (FLUOstar Omega Microplate Reader, BMG LABTECH, Germany). DMAPs were visualized and imaged using a stereomicroscope (Leica EZ4 D, Leica Microsystems, Milton Keynes, UK). Constant compressive force was applied through a TA-TX2 Texture Analyser (Stable Microsystems, UK). Optical coherence tomography (OCT) was carried out in an EX-101 device (Michelson Diagnostics Ltd., Kent, UK). Fluorescence microscopy was carried out on an EVOS FL microscope (Thermo Fisher Scientific, USA). Two-photon fluorescence microscopy was carried out in a Leica TCS SP8- MP multiphoton excited fluorescence upright microscope (Leica Microsystems, UK). *In vivo* fluorescence was evaluated with In-Vivo Xtreme equipment (Bruker, Germany). Flow cytometry was carried out in a CytoFLEX cytometer (Beckman Coulter, USA). Confocal microscopy was performed using a Leica SP5 HyD Confocal Microscope (Leica, Germany).

### 2.5 Preparation of MSN-loaded DMAP

MSN-loaded DMAP was prepared using a negative silicone mold with a design containing 600 pyramidal microneedles (750 μm in length) through the following procedure: i) the microneedle tip region was filled with MSNs in powder form using a spatula; ii) a 20% (w/w) polymer solution (PVA and PVP at a 1:1 weight ratio) in deionized water was added to each mold, followed by centrifugation and removal of excess polymer solution; iii) 800 μL of a 40% (w/w) polymer solution (PVA and PVP at a 1:1 weight ratio) in deionized water was added to each mold, followed by centrifugation; iv) the samples were left to dry for 24 h at room temperature and for 24 additional hours at 37°C. Then, the DMAP was removed from the mold and stored until further use. Control DMAPs without MSNs were prepared by a similar procedure, skipping steps i) and ii).

### 2.6 *In vitro* evaluation of DMAP insertion and dissolution

First, DMAP insertion was evaluated *in vitro* using a Parafilm M® insertion model[36] by applying a compressive force of the DMAP against 8 layers of Parafilm M® for 30 seconds, either 32 N using a Texture Analyser or manually using thumb pressure (32 N was previously selected as a comparable force to the force a human produces when applying thumb pressure on microneedles[36,37]). The depth of insertion was then evaluated by examining the different Parafilm M® layers under a stereomicroscope. DMAP dissolution was then evaluated by introducing DMAP in PBS and imaging the microneedles at different time points using a stereomicroscope. Finally, DMAP insertion and dissolution were also evaluated in a 3% agarose gel. Five minutes after insertion in the agarose gel, the baseplate of the DMAP was removed, and the fate of FITC-labelled or fluorescein sodium-loaded MSNs was evaluated 1 h later (after incubation at 37°C) using fluorescence microscopy.

### 2.7 *Ex vivo* experiments using neonatal porcine skin

Full thickness neonatal porcine skin was used as a skin model, with samples obtained from stillborn piglets and immediately (<24 h after birth) excised. Skin samples were shaved and stored in sealed Petri dishes at −20°C until use. Prior to use, skin samples were equilibrated in PBS. Insertion of DMAP into neonatal porcine skin was carried out as described for the Parafilm M® *in vitro* model. Insertion was evaluated by OCT, and dissolution was evaluated after different time points at 37°C. During DMAP *in situ* dissolution, samples were kept in a sealed container where PBS-wetted paper was placed below the skin samples to prevent them from drying. Quantification of deposited fluorescent MSNs was carried out by fluorimetry following the extraction of the MSNs from excised skin into PBS by thorough sonication in an ultrasound bath. The diffusion of FITC-labelled MSNs or fluorescein sodium (from loaded MSNs) from DMAP across neonatal porcine skin was evaluated using Franz diffusion cells (Crown Glass Co. Inc., Sommerville, USA). Receptor compartments were filled with PBS, and the temperature was controlled during the experiment at 37°C. Skin samples were secured to the donor compartment of the diffusion cell using cyanoacrylate glue with the *stratum corneum* side facing the donor compartment. DMAPs were inserted as previously described into the center of the skin sample. DMAPs were kept in place during the experiment by a cylindrical metal weight (diameter 11 mm, mass 11.5 g) on their upper surface. With DMAP in place, donor compartments were mounted onto the receptor compartments of the Franz cells. Using a long needle, 0.2 mL of sample was removed from the receptor compartment at defined time intervals and replaced with an equal volume of PBS. Sink conditions were maintained throughout the experiment. The concentrations of FITC-labelled MSNs or fluorescein sodium in the receiver medium were determined by fluorimetry.

### 2.8 *In vitro* evaluation of MSNs in a model of dendritic cells

A mouse dendritic cell line (DC 2.4) was used to evaluate the immunological effect of MSNs [27,38]. The day prior to the experiment, 50,000 DC2.4 cells were seeded in each well, using a 96 well plate. For cellular uptake experiments, DC 2.4 cells were incubated with RITC-labelled MSNs for 2 h at a concentration of 10 μg/mL (in complete culture medium, RPMI-1640 supplemented with 10% fetal bovine serum, nonessential amino acids, L-glutamine and β-mercaptoethanol, as recommended by the distributor (Sigma−Aldrich)) at 37°C and 5% CO_2_. Twenty-four hours later, nanoparticle uptake was evaluated by flow cytometry and fluorescence microscopy. To evaluate the biological effect, changes in the expression of CD40 (a marker of dendritic cell activation) were assessed by flow cytometry after incubation with empty and OVA- loaded nanoparticles (nonlabelled).

### 2.9 *In vivo* experiments in mice

Mouse studies were carried out following Spanish national and European regulations (RD1201/2005, 32/2007, 2010/63/UE and RD53/2013). The mice were hosted at IBIMA-Plataforma BIONAND (Registration No. ES 290670001687). All procedures followed the 3R principles and received appropriate regulatory approval before starting (protocol 18/11/2021/180 approved by both the Institutional Ethics Committee and by *Consejería de Agricultura, Ganadería, Pesca y Desarrollo Sostenible, Junta de Andalucía*). The mice were anaesthetized during the different procedures and finally sacrificed by cervical dislocation. After obtaining the corresponding samples, the mice were stored and incinerated according to institutional guidelines.

To evaluate DMAP insertion and MSN deposition in mice, DMAP loaded with FITC-labelled M- MSNs was used. Five- to six-week-old BALB/c mice (both male and female, n=3 mice per group) from Janvier Lab (Saint-Berthevin Cedex, France) were used. For DMAP administration, back hair was first removed from the mice by using a hair-removal cream under intraperitoneal anaesthesia (xylazine + ketamine mixture). After washing the skin with saline solution to remove the cream and gently drying it with paper, the DMAP was inserted on the back skin by applying appropriate pressure for 30 seconds with the mice still under anaesthesia, and then adhesive tape was used to fix the DMAP in the same position. After 2 h, the baseplates were removed, and excess polymer on the skin surface (not inserted) was removed with PBS. After different time points, nanoparticle fluorescence in the mice was examined using an *in vivo* imaging system. At the endpoint (3 days after DMAP administration), the mice were euthanized by cervical dislocation under anaesthesia, and the skin was observed by fluorescence microscopy. The deposited MSN amount was quantified as described for the *ex vivo* experiments. Finally, tissue sections from skin were obtained and evaluated under fluorescence confocal microscopy after immunofluorescence staining for immune cells (using primary antibodies against F4/80 for murine macrophages and CD11c for mouse dendritic cells: rabbit anti-mouse CD11c primary antibody and goat anti-mouse F4/80 primary antibody; and fluorescent secondary antibodies for visualization: chicken anti-rat IgG (H+L) cross-adsorbed secondary antibody-Alexa Fluor™ 647; goat anti-rabbit IgG (H+L) cross-adsorbed secondary antibody-Alexa Fluor™ 555).

To evaluate the immunological effect of antigen-loaded MSNs administered through DMAP, OVA-loaded XL-MSNs were administered either subcutaneously or in DMAP once a week for 3 weeks. Five- to six-week-old BALB/c mice (male, n=3 mice per group) from Janvier Lab (Saint-Berthevin Cedex, France) were used. One week after the last administration, the mice were anaesthetized intraperitoneally, and blood was obtained from the retroorbital plexus before euthanizing the animals. Different specific anti-OVA antibodies (IgG1, IgG2a, IgG2b and IgE) were determined from the mouse sera by ELISA using biotinylated rat anti-mouse antibodies. For the ELISAs, high binding ELISA 96-well plates were coated with OVA. After blocking the plate with a casein- containing buffer solution, the mouse sera were added (1:8 dilution for IgE detection, 1:50 dilution for IgG detection) and incubated overnight at 4°C. Then, biotinylated secondary antibodies were added, followed by the addition of avidin-horseradish peroxidase (HRP). Finally, the results were obtained by measuring the colorimetric conversion of an HRP substrate (TMB) after stopping the reaction with H_2_SO_4_ using a plate reader (λ_ABS_=450 nm). Thorough washing of the plate with PBS containing 0.05% Tween 20 was carried out between the different steps in the protocol.

## 3. Results and Discussion

### 3.1 Synthesis and characterization of MSNs

MSNs with different pore sizes (from smaller to larger: S-MSN, M-MSN, L-MSN and XL-MSN) were prepared and characterized. The size histograms obtained by DLS show peak particle diameters of approximately 100 nm (**Figure 1A-D**), with Z average values in the range of 110-160 nm and narrow size distributions (polydispersity index, PDI, below 0.2) for all the obtained particles (**Table 1**). The nanoparticle surface charge was negative for all the prepared MSNs, as would be expected by the presence of silanol groups on the external surface of silica nanoparticles (**Table 1**). The round morphology and porosity of the MSNs could also be observed in the TEM micrographs of the prepared nanoparticles (**Figure 1E-H**).

**Table 1.** Characterization of the prepared MSNs by DLS, Z potential and N_2_ adsorption.

|  | S-MSN | M-MSN | L-MSN | XL-MSN |
| --- | --- | --- | --- | --- |
| Hydrodynamic diameter (Z Average, nm) | 123.5 ± 2.33 | 130.2 ± 0.89 | 117.1 ± 2.39 | 155.4 ± 2.08 |
| PDI | 0.15 ± 0.02 | 0.19 ± 0.01 | 0.12 ± 0.03 | 0.16 ± 0.03 |
| Z Potential (mV) | -8.31 ± 0.7 | -24.5 ± 0.2 | -12.1 ± 0.4 | -32.8 ± 0.4 |
| BET* Surface Area (m<sup>2</sup>/g) | 321.9 ± 24.6 | 522.8 ± 148.9 | 560.9 ± 149.3 | 679.4 ± 11.1 |
| Pore volume (cm<sup>3</sup>/g) | 0.39 ± 0.02 | 0.71 ± 0.21 | 0.93 ± 0.18 | 1.38 ± 0.05 |
| Pore width (nm) | 3.33 ± 0.21 | 5.65 ± 0.14 | 8.46 ± 0.09 | 11.77 ± 0.64 |

\*Brunauer–Emmett–Teller (BET) method

**Figure 1.**
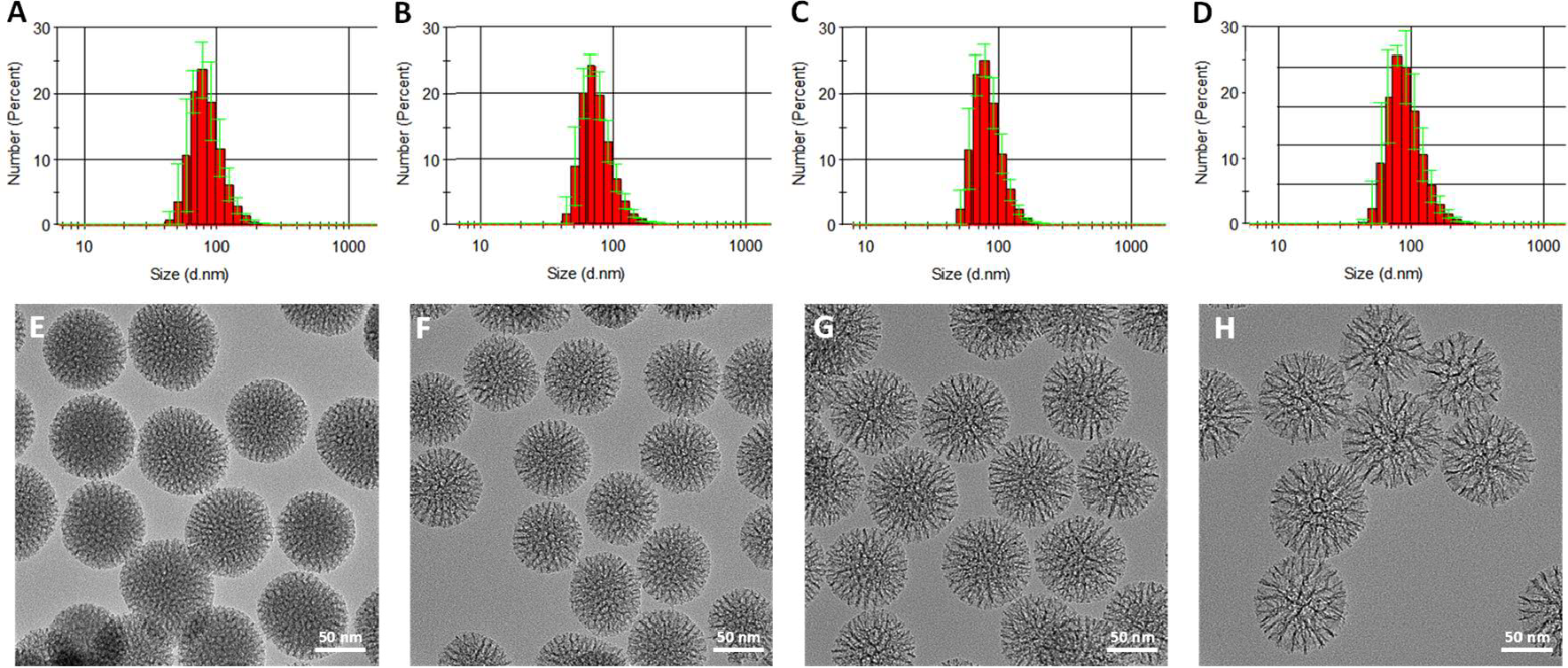
DLS size distribution histograms of MSNs: S-MSN (A), M-MSN (B), L-MSN (C) and XL- MSN (D); TEM micrographs of MSNs: S-MSN (E), M-MSN (F), L-MSN (G) and XL-MSN (H).

The textural properties of the prepared MSNs were further confirmed by N_2_ adsorption (**Table 1**). All the materials presented large surface areas (in the range of 300-700 m^2^/g), typical of mesoporous silica materials. The pore diameters obtained by N_2_ adsorption confirm the successful preparation of MSNs with 4 different pore sizes, in the range of 3 to 12 nm (S-MSN<M- MSN<L-MSN<XL-MSN). These results were in good agreement with the characteristics of MSNs prepared by other authors using the same method[26].

The effect of pore size on cargo loading and release was evaluated using fluorescein sodium salt as a model for small molecule drugs and OVA as a model for therapeutic proteins and macromolecules (**Figure 2**). The loading of OVA was maximum for extralarge pore particles (XL- MSN) and decreased as pore size was reduced. This result was in good agreement with previous reports that have also shown that MSNs with extralarge pore sizes presented increased OVA loading compared to particles with smaller pores[27]. On the other hand, fluorescein sodium salt loading was maximum for particles with small or medium pores (S-MSN and M-MSN) and drastically decreased for particles with larger pores. These data indicate that to obtain optimal cargo loading in MSNs, the pores should be large enough to accommodate the cargo, but if the pore size is too large, the loading efficiency decreases. Thus, the pore size should be tuned depending on the molecular weight of the cargo, requiring smaller pores for small molecules and larger pores for macromolecules such as proteins. Despite this, when cargo release experiments were carried out, larger pore particles presented faster release kinetics for both model cargos, as the larger pores provide easier accessibility to the solvent, which drives release. This result is also in good agreement with previous reports[27]. Furthermore, taking these results into account, a combination therapy scheme could be envisioned where drugs of different molecular weights can be coadministered in a cocktail of MSNs of varying pore sizes, each one of which is optimized for one of the drugs in the combination.

**Figure 2.**
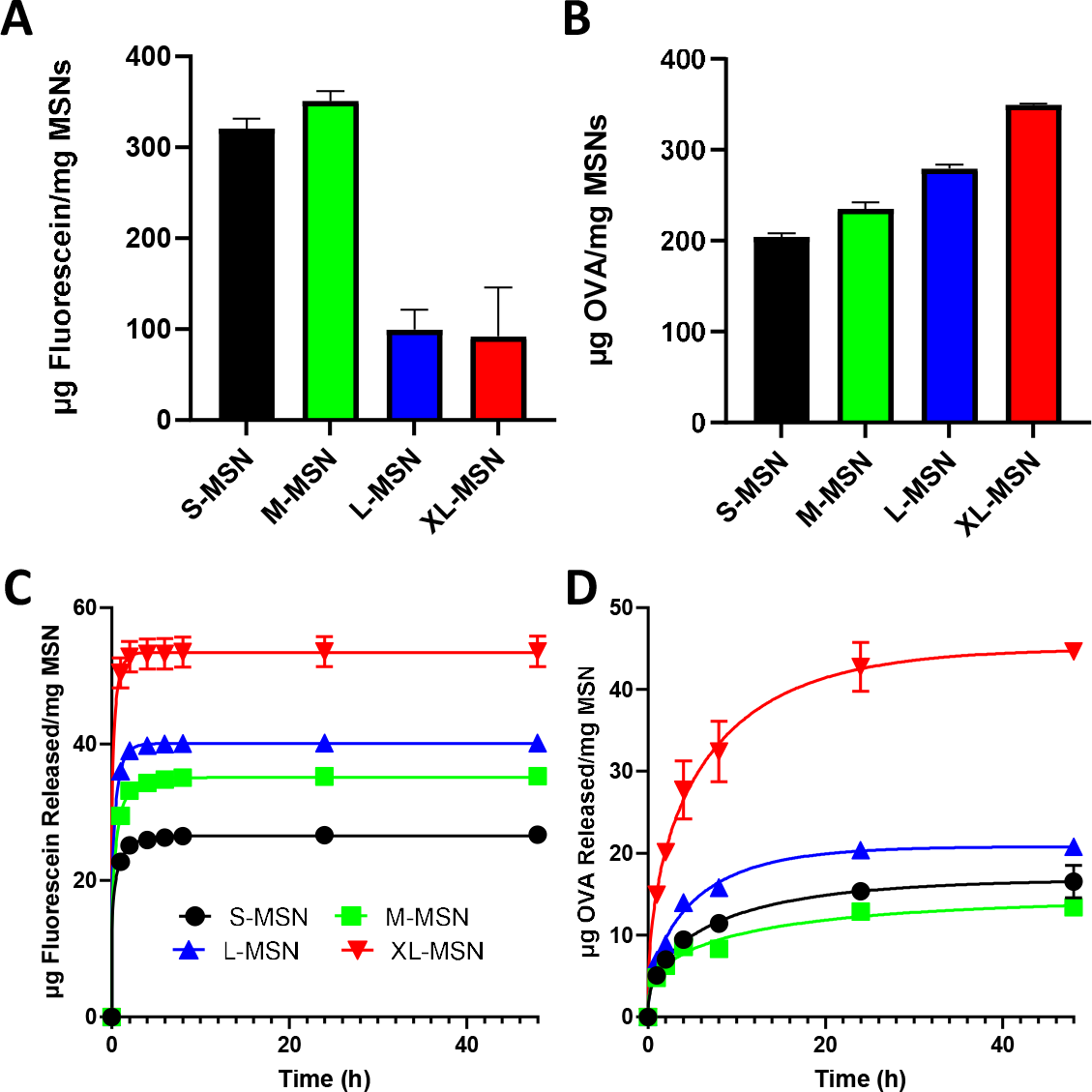
Cargo loading and release experiments in/from MSNs of different pore sizes. Loading of fluorescein sodium salt (A) and OVA (B); Release of fluorescein sodium salt (C) and OVA (D). Data are means ±SDs, n=3.

### 3.2 Preparation and characterization of MSN-loaded DMAP

Then, DMAP-containing MSNs of different pore sizes were prepared using PVP and PVA (2 water-soluble polymers). When attempting to disperse MSNs in a solution of PVP and PVA to prepare DMAP, we found that the maximum weight % of MSNs that could be added in the mixture that still enabled filling up the molds used to prepare DMAP was 30%. When larger percentages of MSNs were included in the polymer mixture, its viscosity was too high to properly fill the mold used to obtain DMAP. As our aim was to prepare DMAP with a large amount of MSNs located in the microneedle tips, we developed an alternative method by first filling the mold with MSNs in powder form and later adding the polymers in solution. After following this process, the stereomicroscopy images of the obtained DMAP (**Figure 3**) confirm the successful preparation of the desired arrays presenting well-defined microneedles of the expected length (*ca*. 750 μm) and with intact tips. No morphological differences were found between the blank DMAP (without nanoparticles) and those containing the different types of MSNs. Furthermore, the SEM micrographs (**Figure 3P-Y**) confirm the presence of MSNs, which make up most of the microneedle tips, as seen in the higher magnification micrographs (**Figure 3U-Y**). These results confirm the preparation of DMAP in which the majority of the microneedle tips are composed of MSNs, in contrast with previous reports, where the amount of MSNs included in DMAP formulations was relatively low [24]. Furthermore, as only the microneedle tips will be inserted inside the skin upon DMAP application, the MSNs should be selectively located in the microneedle tips to avoid unnecessary wastage of drug-loaded particles in future therapeutic applications of these formulations. To confirm the location of the MSNs within the patches, FITC-labelled M-MSNs were prepared and used to obtain DMAP. The stereomicroscopy and two-photon fluorescence microscopy images obtained from these formulations (Figure S1) confirm the presence of the MSNs only in the microneedle tips.

**Figure 3.**
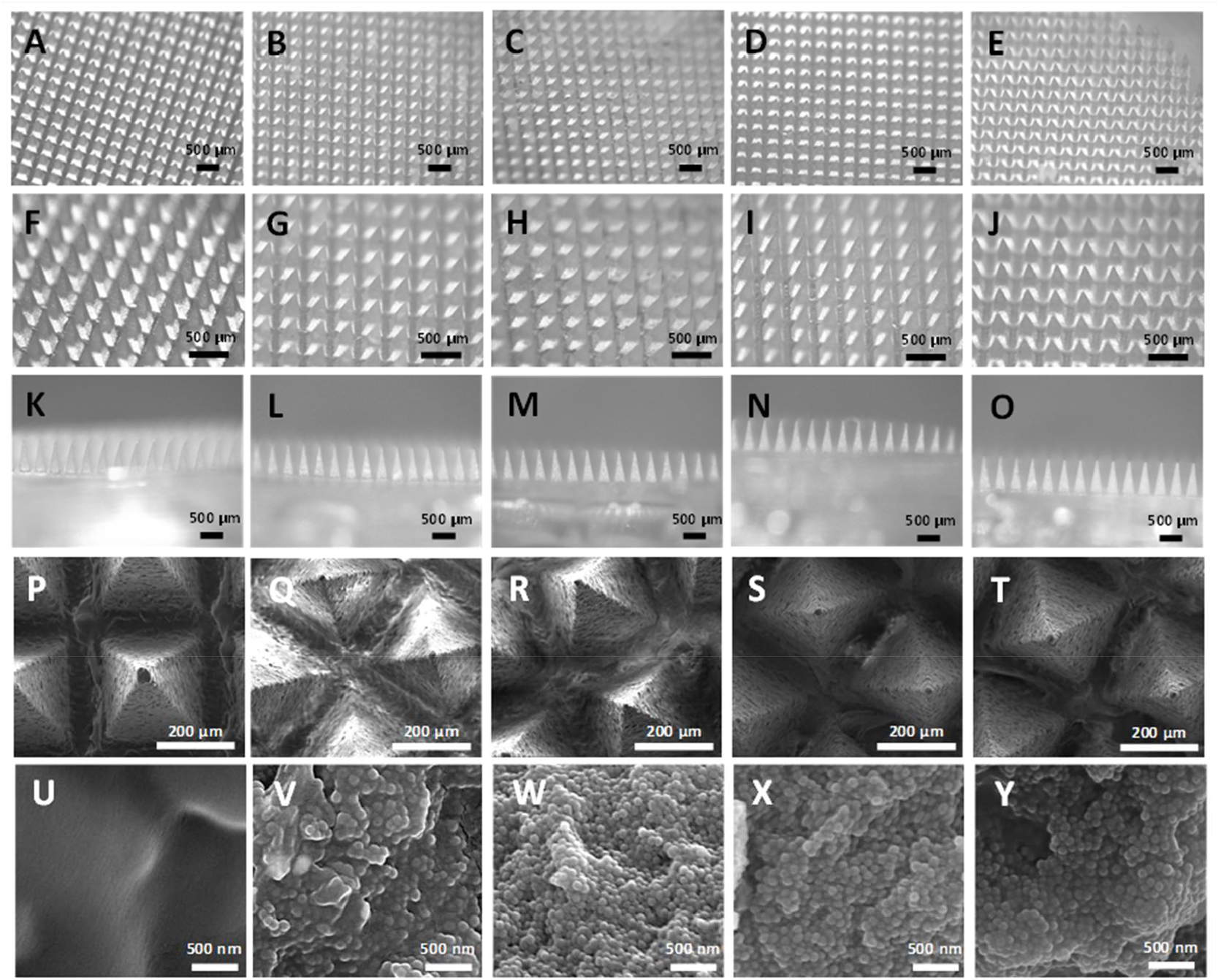
Stereomicroscopy (A-O) and SEM (P-Y) images of DMAP-containing MSNs at different magnifications and orientations. Blank DMAP without MSN (A,F,K,P,U); S-MSN (B,G,L,Q,V); M- MSN (C,H,M,R,W); L-MSN (D,I,N,S,X) and XL-MSN (E,J,O,T,Y).

### 3.3 *In vitro* and *ex vivo* evaluation of MSN-loaded DMAP

The mechanical properties of the prepared DMAP and whether they allow for their insertion in skin were first evaluated *in vitro* using a previously reported Parafilm M® model[36]. DMAP insertion was evaluated either using a Texture Analyser applying 32 N for 30 seconds or by manually applying thumb pressure for 30 seconds. The results (**Figure 4**) show that the insertion of all the prepared DMAP was similar when inserted in the same way, regardless of whether the DMAP contained any kind of MSN. Furthermore, the DMAPs were inserted more efficiently by manual application (successfully piercing through the 3^rd^ Parafilm M® layer, with a depth of 378 μm) compared to the ones inserted using the Texture Analyser equipment (which only reached the 2^nd^ layer, at a depth of 252 μm). These results are in good agreement with previous reports of microneedle patches that show similar insertion in the Parafilm M® model, as well as deeper insertion upon manual force [39]. Finally, the mechanical properties of the prepared DMAP were also proven to be adequate for further evaluation, as there was no significant change (p>0.54) in microneedle length after insertion in the Parafilm M® model, neither by manual nor Texture Analyser application (**Figure 4B**). The *in vitro* dissolution of the different DMAP formulations in aqueous media was also confirmed by immersing them in PBS. Three minutes after immersion, the microneedles of all DMAP were fully dissolved, as observed by stereomicroscopy (**Figure S2**). By dissolving in PBS DMAP containing FITC-labelled M-MSNs and then measuring the fluorescence of the resulting suspensions, the amount of nanoparticles loaded in each DMAP could be estimated to be 2.33 ± 0.04 mg M-MSNs/DMAP.

**Figure 4.**
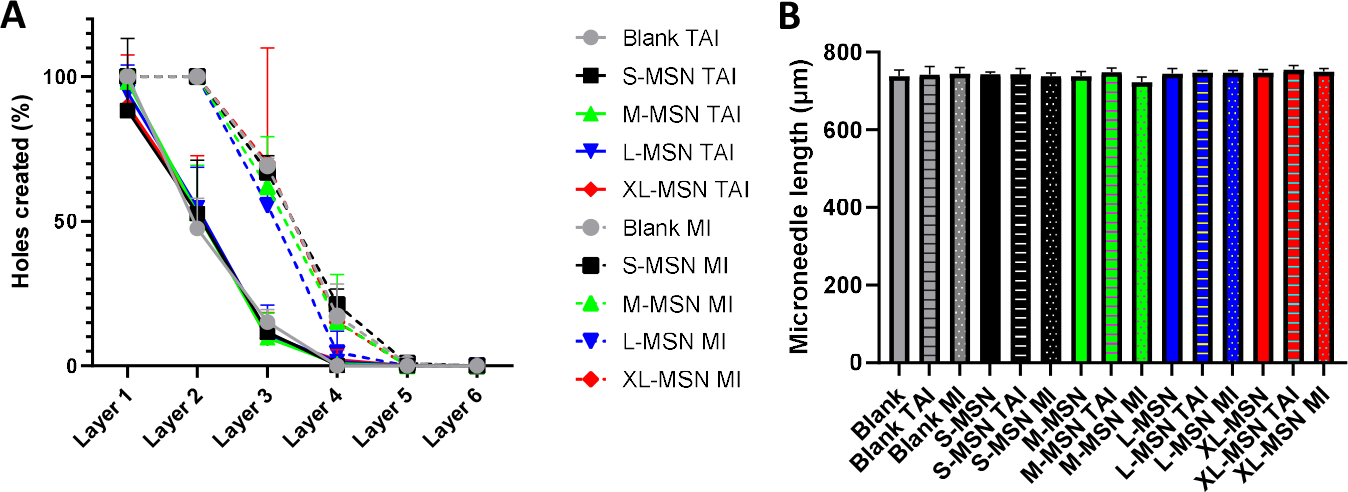
DMAP insertion in the Parafilm M® model either using a Texture Analyser (TAI) (continuous lines) or through manual insertion (MI) (dotted lines) (A). Blank (not containing MSNs) or MSN-containing DMAP were evaluated. Microneedle lengths before and after insertion in the Parafilm M® model (B). Data are means ±SDs, n=3.

The next step was to evaluate the insertion and dissolution of DMAP in neonatal porcine skin, which will provide more relevant information towards the potential use of these formulations in humans. After manual *ex vivo* application in porcine skin, the OCT images confirm the successful insertion of the microneedles inside the porcine skin, with an average insertion length of 472 ± 26.7 μm (62.9 ± 3.5% of the total microneedle length) (**Figure 5**). Furthermore, the successful dissolution of the microneedles inserted in the skin was also observed, with partial dissolution taking place in DMAP inserted for 30 min at 37°C and reaching almost complete dissolution at 60 min (Figure 5B-C). At this time (60 min), the amount of FITC-labelled M-MSNs that had been deposited in the skin was 0.75 ± 0.02 mg, which was 20.9 ± 7.26% of the total amount present in the DMAP (Figure 5D-F).

**Figure 5.**
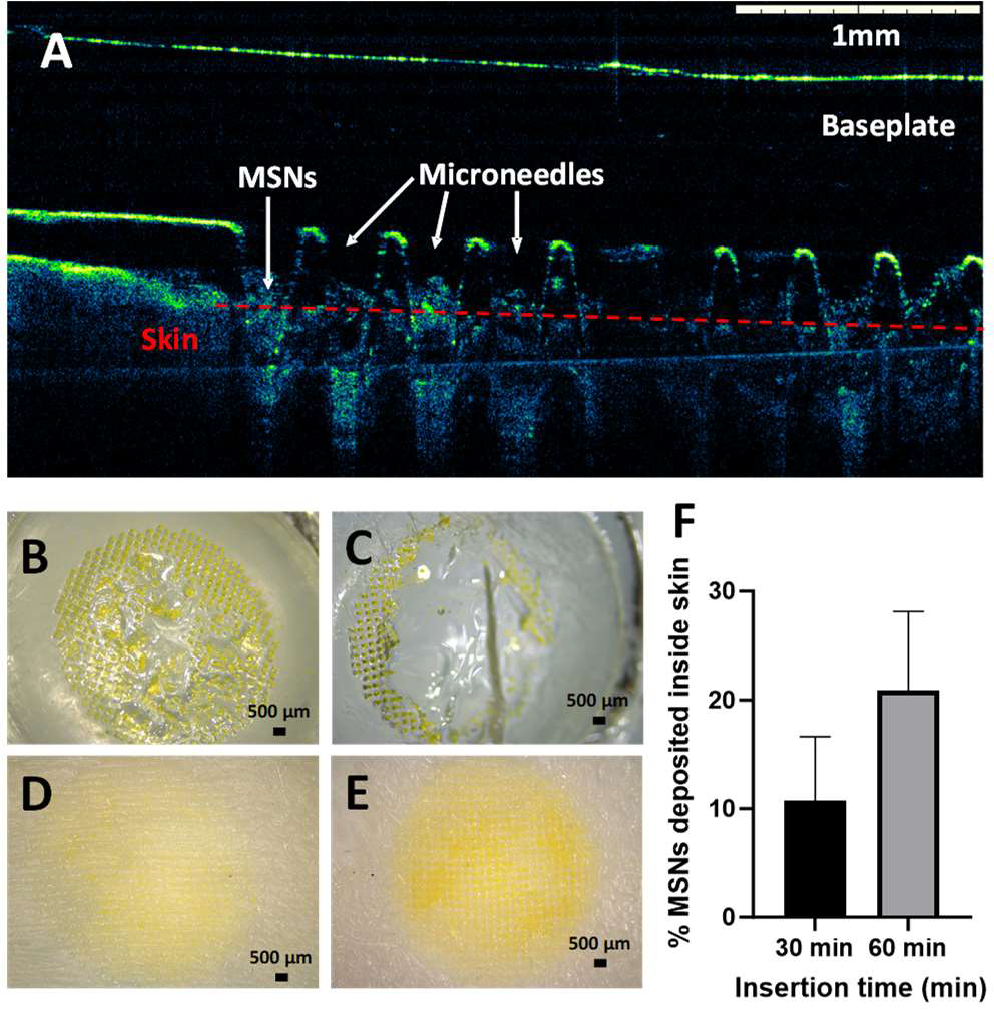
DMAP insertion and dissolution *ex vivo* using neonatal porcine skin. OCT image of M- MSN-containing DMAP inserted in neonatal porcine skin (A). Photographs of FITC-labelled M- MSN-containing DMAP after insertion in skin for 30 (B) or 60 (C) min. Photographs of neonatal porcine skin after removal of FITC-labelled M-MSN-containing DMAP that had been inserted for 30 (D) or 60 (E) min. Quantification of FITC-labelled M-MSN deposited inside neonatal porcine skin at different insertion times (F). Data are means ±SDs, n=3.

As seen in the results shown in **Figure 2**, cargo loading and release in MSNs is strongly affected by the interaction between cargo molecular weight and the pore size of the nanoparticles. For this reason, we postulate the potential use of combinations of MSNs of various pore sizes for combined drug codelivery, where each type of MSN is optimized for one drug in the mix. With this goal in mind, the next step was to evaluate whether the method developed to prepare DMAP would enable patches with combinations of different MSNs to be obtained. To evaluate this, we selected MSNs that presented optimal loading for each of the model molecules previously used: M-MSNs (which presented the largest fluorescein sodium salt loading capacity) and XL-MSNs (which presented the largest OVA loading capacity). To visualize each type of particle independently, we prepared and used differently labelled MSNs: FITC-labelled M-MSNs and RITC-labelled XL-MSNs. After mixing (in suspension) both types of MSNs in a 1:1 weight ratio and drying the combination, the MSN mix was used to prepare DMAP following the same method as previously described. The characterization of these new DMAPs confirms the successful preparation of DMAPs containing a combination of MSNs of different pore sizes (**Figure 6, Figure S3, Table S1**). Furthermore, DMAP containing a combination of FITC-labelled M-MSNs and RITC- labelled XL-MSNs presented analogous insertion and nanoparticle deposition in neonatal porcine skin as those previously evaluated containing only one MSN type (Figure S4, Table S2).

**Figure 6.**
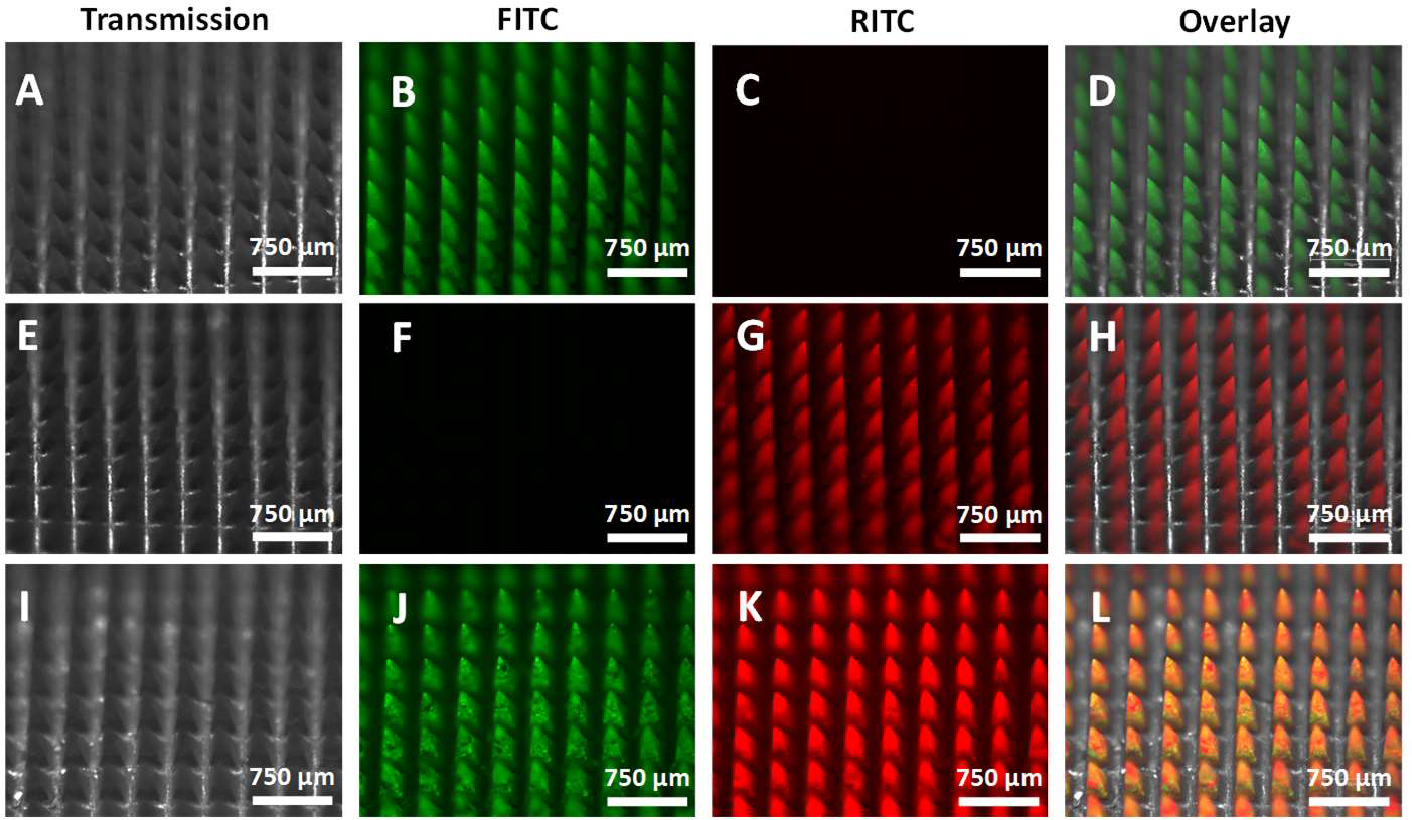
Fluorescence microscopy images of DMAP containing FITC-labelled M-MSNs (A-D), RITC-labelled XL-MSNs (E-H) or a combination of both types of MSNs (I-L).

In a potential future application of these DMAPs, after insertion and microneedle dissolution, drug-loaded MSNs will be deposited inside the skin. The fate of both MSNs and their cargo after deposition in the skin should therefore also be evaluated to determine the potential of these formulations. To evaluate the diffusion of both the nanoparticles and their cargo, DMAP with 2 different nanoparticle systems was prepared: FITC-labelled M-MSNs (to analyse the fate of the nanoparticles) and fluorescein sodium salt-loaded M-MSNs (to analyse the fate of the fluorescent cargo as a model for a drug being released from the nanoparticles). In the first experiment, an agarose gel was used as a model for skin tissue. After insertion of DMAP, the microneedles dissolve and leave the MSNs inside the agarose gel. One hour later, sections of the gel were cut and evaluated by fluorescence microscopy. The results (**Figure S5**) show that while the MSNs remain in the site of deposition, the cargo (fluorescein sodium salt) diffuses out of the nanoparticles and into the gel. This would indicate that the nanoparticles would remain in the site of insertion, acting as a depot system and providing a sustained release of the drug to the surrounding skin tissue. To confirm these results, a second experiment was carried out in a Franz cell setup. The same DMAP as those from the previous experiment (plus an additional control DMAP with nonencapsulated fluorescein sodium salt) were inserted in neonatal porcine skin that acted as the membrane separating the donor and receptor compartments. At different time points, samples were taken from the receptor compartment, and the amount of FITC-labelled M-MSNs or free fluorescein sodium salt was evaluated by fluorimetry. The results (**Figure 7**) confirm those from the previous experiment, showing that MSNs are not capable of diffusing through the skin tissue, as no nanoparticle fluorescence was detected in the receptor compartment. On the other hand, fluorescein sodium salt crosses through the skin tissue after being deposited with DMAP. Importantly, encapsulation of the dye inside M-MSNs provided sustained release compared to the use of DMAP with a nonencapsulated fluorophore. Taken together, these results indicate that the MSNs deposited in the skin through DMAP remain at the site of insertion and act as a drug reservoir. Then, the sustained release of the drug would provide a continuous flow of the drug towards both the surrounding tissue or even systemic circulation (depending on the release kinetics and dose of the drug or nanoparticles included in the patches).

**Figure 7.**
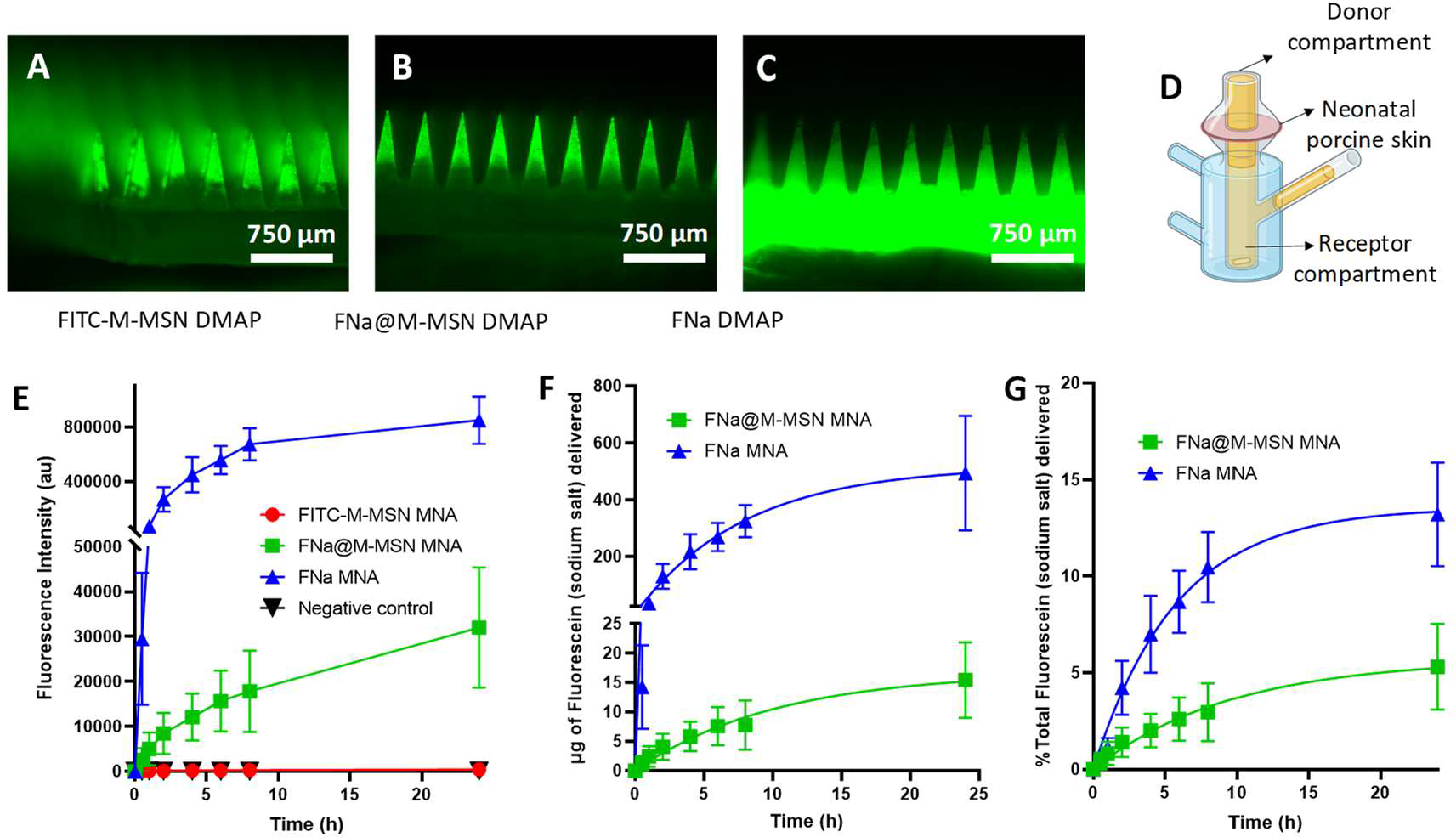
Franz cell experiment with neonatal porcine skin. Fluorescence microscopy images of the different DMAPs used in the experiment: FITC-labelled M-MSNs (A), fluorescein sodium salt-loaded M-MSNs (B), and nonencapsulated fluorescein sodium salt (C). Schematic representation of the Franz cell setup (D). Fluorescence intensity in the receptor compartment (indicating transdermal delivery) at different time points after DMAP insertion in neonatal porcine skin (E). Amount (in μg) of total fluorescein sodium salt delivered transdermally at different time points (F). Percentage of total fluorescein sodium salt delivered transdermally at different time points (G). Data are means ±SDs, n=5.

### 3.4 *In vivo* evaluation of MSN-loaded DMAP

Once the characteristics and performance of MSN-loaded DMAP had been evaluated through the different *in vitro* and *ex vivo* experiments described above, a series of *in vivo* tests were carried out using a mouse model to examine the therapeutic potential of the developed platform. First, the *in vivo* insertion, microneedle dissolution and nanoparticle deposition were evaluated by a combination of *in vivo* fluorescence and *ex vivo* fluorescence stereomicroscopy (**Figure 8**). The results confirm the successful insertion, microneedle dissolution and nanoparticle deposition, as nanoparticle fluorescence is clearly visible in both the *in vivo* fluorescence imaging and the *ex vivo* fluorescence stereomicroscopy images 3 days after DMAP administration to the mice. The insertion of the DMAP was successful in both male and female mice, requiring 2 hours of insertion for almost complete microneedle dissolution in the skin (with only partial dissolution after 1 h). In the *ex vivo* fluorescence stereomicroscopy images, the microneedle pattern is clearly visible in the green fluorescence, confirming successful nanoparticle deposition in DMAP- treated mice (both male and female), while no fluorescence was observed in the skin of control animals. The quantification of the total amount of deposited FITC-labelled M-MSNs (carried out using the same method as described for the *ex vivo* experiments in neonatal porcine skin) provided an estimation of 134.2 ± 79.3 μg of MSNs deposited inside the mouse skin per DMAP application.

**Figure 8.**
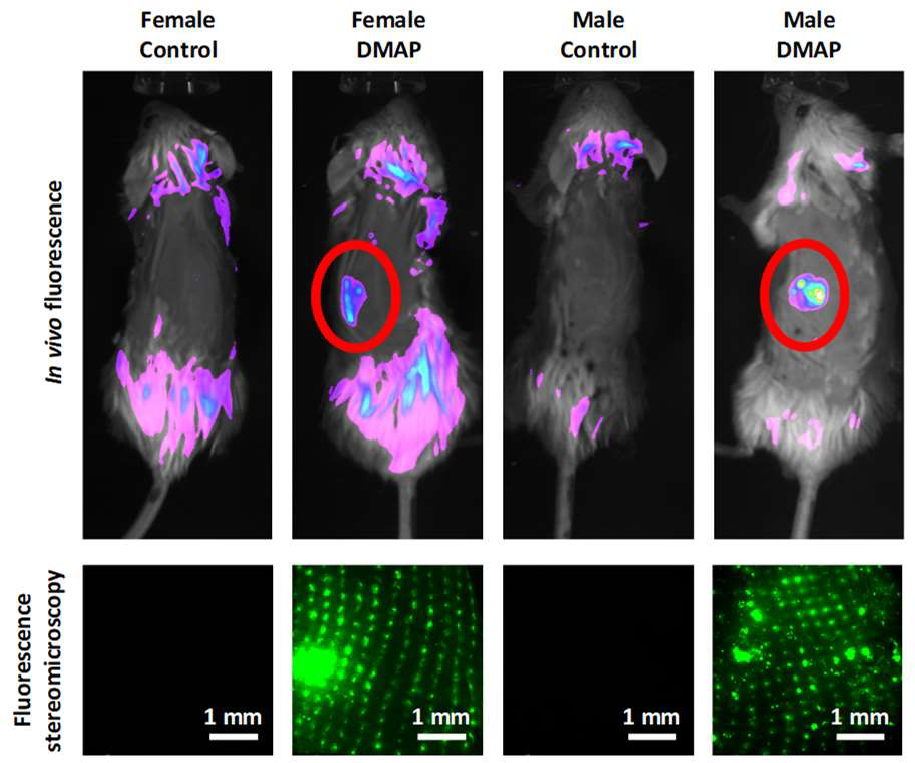
DMAP insertion, dissolution and MSN deposition *in vivo. In vivo* fluorescence imaging of both controls and FITC-labelled M-MSN-loaded DMAP-inserted male and female mice taken 3 days after DMAP administration (top). *Ex vivo* fluorescence stereomicroscopy of skin excised from the same mice showing fluorescence of the deposited FITC-labelled M-MSNs (bottom).

Using the DMAP-inserted skin samples from the previous experiment, the fate of the deposited nanoparticles was also evaluated by confocal microscopy using immunofluorescent labelling. In previous *in vitro* and *ex vivo* experiments, we observed that deposited MSNs remain in the site of insertion without significant diffusion into the surrounding skin tissue. However, these experiments were performed in the absence of living cells in the skin tissue that might engulf the nanoparticles and determine their final fate and biological performance. Two different types of fluorescent labels were used: F4/80 staining (a traditional marker for macrophages[40,41]) and CD11c staining (a traditional marker for dendritic cells)40,42]). In agreement with previous reports, we found (**Figure 9**) that the majority of the dermal cells in mouse skin express F4/80, while a much smaller percentage express CD11c [40]. In any case, most of the dermal cells are antigen-presenting cells, either F4/80^+^ or CD11c^+^. Green fluorescence from the nanoparticles was found in all tissue sections around cell nuclei (stained with DAPI), indicating that the nanoparticles are taken up by dermal cells after deposition in the skin from DMAP. Nanoparticle fluorescence was observed inside antigen-presenting cells (which were also positive for either F4/80 or CD11c, **Figure 9E, F, K, L**). As the MSNs deposited in the skin through DMAP were then taken up by antigen presenting cells, a potential application of these systems would be in vaccination or immunotherapy as carriers of different antigens to the target immune cells in the skin. To evaluate the biological effect of antigen-loaded MSNs in antigen-presenting cells, an *in vitro* experiment was first carried out using a mouse cell line commonly employed as a model for dendritic cells (DC2.4[27]). The results (Figure S6) show that DC2.4 cells efficiently uptake MSNs, with a clear pore size effect, as XL-MSNs presented a lower uptake % compared to all other types of MSNs. When the activation of DC2.4 cells was evaluated (by quantifying the % of cells expressing CD40 by flow cytometry), the results showed that OVA-loaded XL-MSNs induced the largest degree of dendritic cell activation, decreasing the % of CD40^+^ cells as the nanoparticle pore size decreased. Furthermore, treatment with free OVA in the absence of a nanocarrier, even at high concentrations (up to 4 μg/mL), did not induce significant cell activation. These results are in good agreement with previous reports that showed that MSN pore size regulates their antigen delivery efficiency[27], with extralarge pore MSNs loaded with OVA (similar to the ones prepared in this work) being the optimal antigen delivery system.

**Figure 9.**
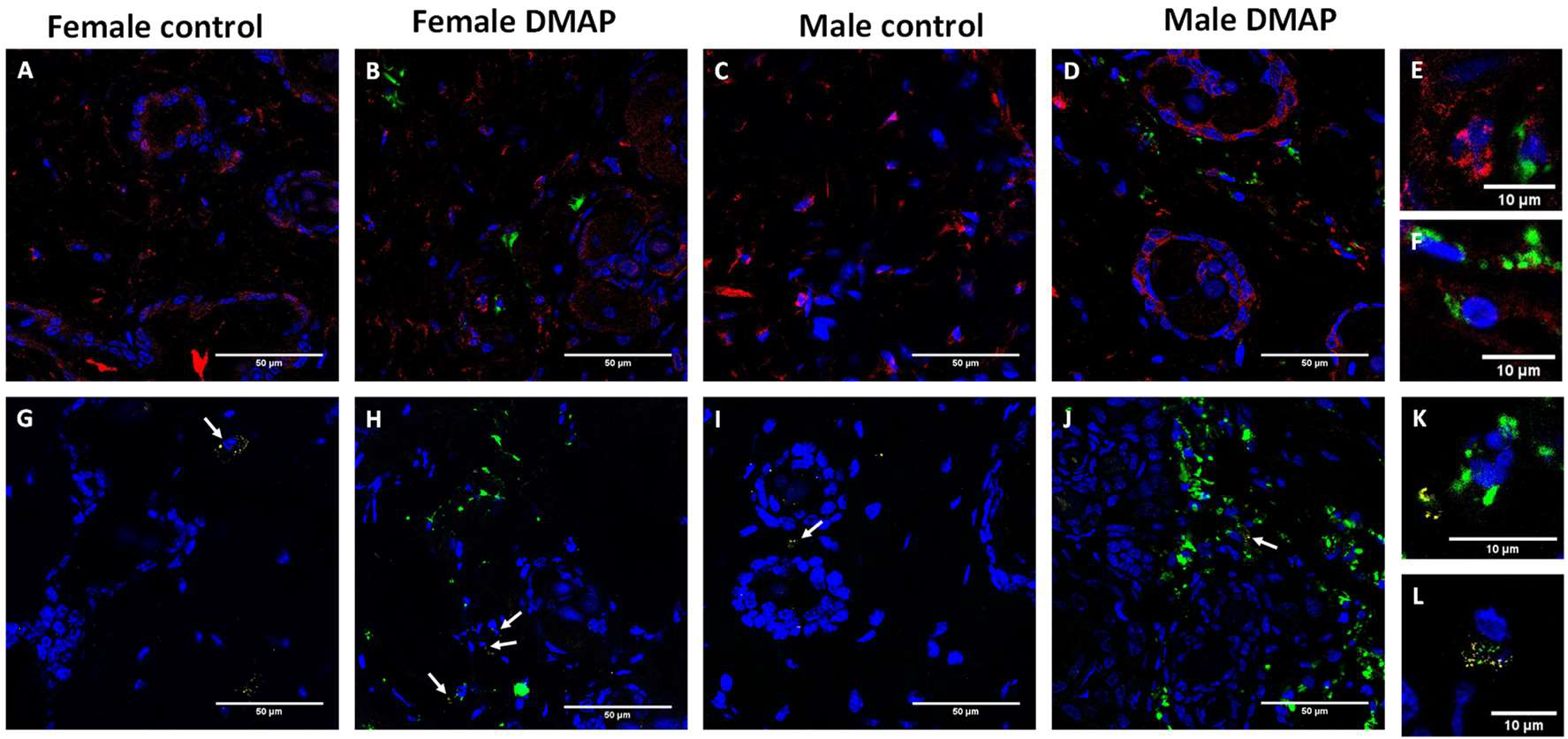
Confocal microscopy images of immunofluorescently labelled mouse skin sections 3 days after FITC-labelled M-MSN-loaded DMAP administration. Female (A,B,G,H) and male (C,D,I,J) mice were used, either as controls (A,C,G,I) or with DMAP application (B,D,H,J). Inserts at higher magnification showing double-positive cells: MSN^+^F4/80^+^ (E, F) or MSN^+^CD11c^+^ (K, L). Blue signal represents DAPI nuclei staining, green signal represents MSN fluorescence, red signal represents F4/80 staining (A-F) and yellow signal (highlighted with white arrows) represents CD11c staining (G-L).

In previous reports, the good potential of XL-MSNs as antigen delivery systems[27,43] was evaluated by subcutaneous injection in mice. While subcutaneous injection is known to produce a strong immune response, the need for trained personnel and the pain associated with the injection could be improved with a needle-free administration system such as the one proposed here. In some applications, there are also safety concerns associated with subcutaneous administration, such as in allergy immunotherapy, in which an undesired allergic reaction to the treatment can be triggered in subcutaneous immunotherapy (with some reports indicating an improved safety profile of microneedle-assisted allergy immunotherapy[44]). To evaluate the prospects of MSN-loaded DMAP for immunization, OVA-loaded XL-MSNs were administered 3 times (weekly) in BALB/c mice either subcutaneously or in DMAP (equivalent nanoparticle dose in both types of administration). OVA-loaded XL-MSNs were used since despite presenting a lower uptake %, their capacity to activate DC2.4 cells was higher. One week after the last administration, blood was obtained, and OVA-specific antibodies were evaluated in sera by ELISA. The results (**Figure 10**) show comparable humoral responses between subcutaneous and DMAP-mediated administration of OVA-loaded XL-MSNs, with significant production of anti-OVA IgG1 and IgG2b compared to nontreated control mice. On the other hand, no significant differences were found between control and nanoparticle-treated mice in the production of specific anti-OVA IgG2a or IgE. These results highlight the promising nature of the developed platform for vaccination or immunotherapy through needle-free administration. Future work will explore the combination of MSNs carrying an immunomodulatory adjuvant with antigen- carrying MSNs for different therapeutic applications, as this work has demonstrated the possibility of preparing DMAP with combinations of different types of MSNs tailored for various therapeutic molecules of a wide range of molecular weights.

**Figure 10.**
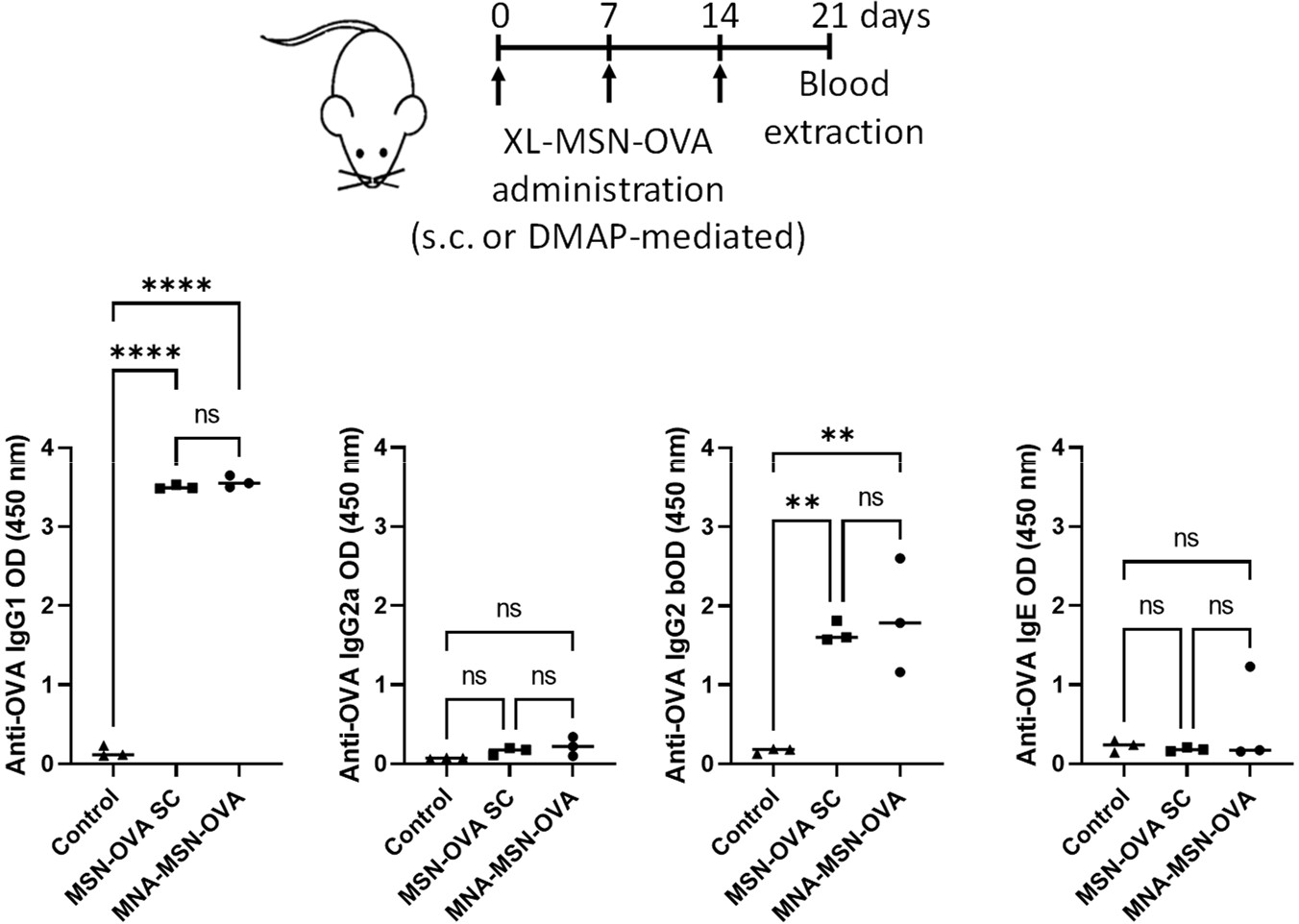
*In vivo* immunization experiments in mice employing OVA-loaded MSNs either by subcutaneous injection or by administration through DMAP. OVA-specific antibodies (IgG1, IgG2a, IgG2b and IgE) were measured in sera by ELISA. Data are means ±SDs, n=3. Statistical analysis by one-way ANOVA. ns p>0.05; ^**^p<0.01; ^****^p<0.001.

## Conclusions

In this work, dissolving microneedle array patches containing large amounts of mesoporous silica nanoparticles of different pore sizes were prepared and characterized. The developed method also enabled the preparation of microneedle array patches containing a combination of different mesoporous silica nanoparticles, which could be useful for the development of combination therapies codelivering different therapeutic cargos. The successful insertion, dissolution and nanoparticle deposition from the microneedles was confirmed through a series of *in vitro, ex vivo* and *in vivo* experiments. As the microneedle-delivered mesoporous silica nanoparticles were found to end up inside antigen presenting cells in the skin tissue, a particularly promising application of these systems would be in vaccination or immunotherapy applications. To confirm this potential application, the immune response to ovalbumin-loaded mesoporous silica nanoparticles in mice was evaluated, showing comparable levels of specific antibody generation after subcutaneous or microneedle-mediated delivery. Based on the promising results presented here, future work will further evaluate the therapeutic potential of this platform for immunotherapy in different disease scenarios, by developing microneedle codelivery systems containing antigenic and adjuvant molecules encapsulated in optimized mesoporous silica nanoparticles.

## Acknowledgements

This work was supported by the Institute of Health “Carlos III” (ISCIII) (PI21/00346), cofounded by the European Union, and by the project PID2022-142781OA-I00 funded by MCIN/AEI/10.13039/501100011033/FEDER, UEU. Funding from Wellcome Trust (grant number WT094085MA) is also gratefully acknowledged. Fellowships from ISCIII were held by JLP (CD19/00250, MV21/00077) and JAC (CD20/00085), with the “Sara Borrell” grant co-funded by the European Social Fund (“El FSE invierte en futuro”). JLP also acknowledges grant RYC2021- 034536-I funded by MCIN/AEI/10.13039/501100011033 and by European Union NextGenerationEU/PRTR. CM holds a “Nicolas Monardes” research contract by Andalusian Regional Ministry Health (RC0004-2021). TEM, confocal microscopy and *in vivo* imaging experiments were performed in the ICTS “NANBIOSIS,” more specifically in the U28 Unit at IBIMA Plataforma BIONAND.

