## Supporting Information for "Dissolving microneedle array patches containing mesoporous silica nanoparticles of different pore sizes as a tunable sustained release platform"

FOR

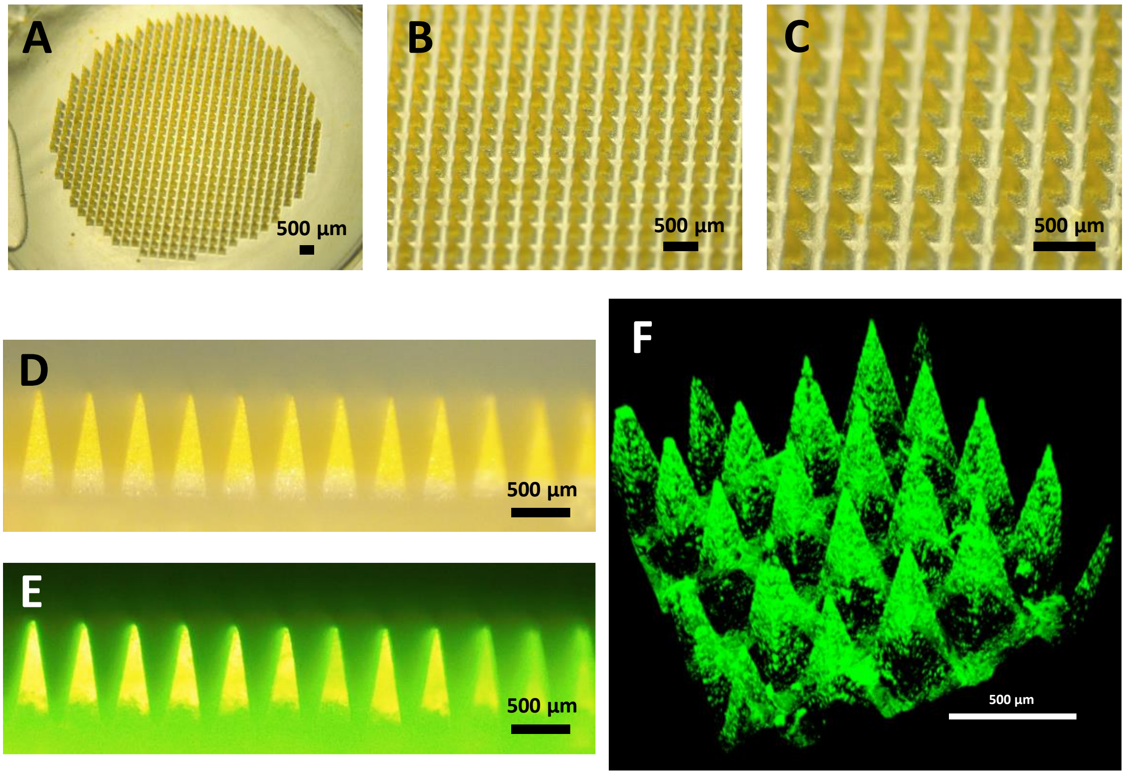


**Figure S1.** DMAP containing FITC-labeled M-MSN. Stereomicroscopy images (A-D), Fluorescence stereomicroscopy (E) and 3D reconstruction of DMAP using Two-photon fluorescence microscopy, all showing selective location of the nanoparticles in the microneedle tips.

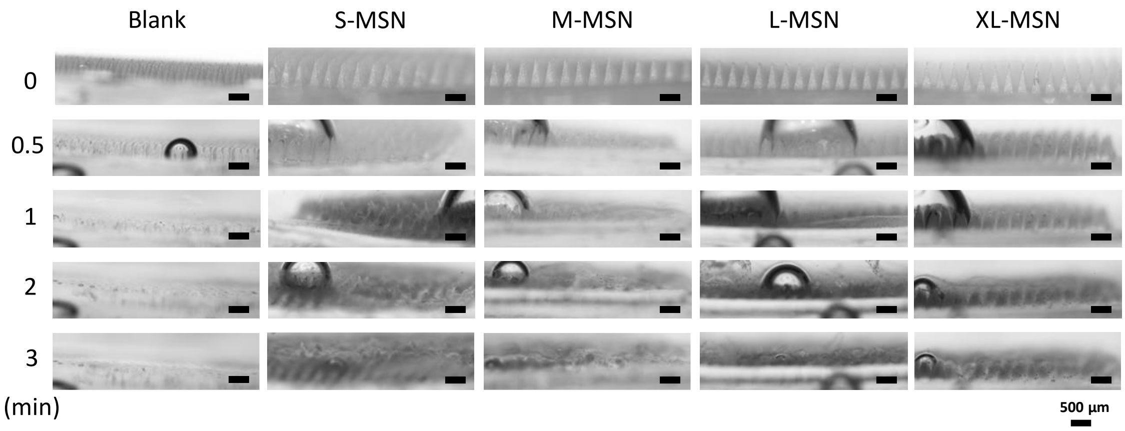


**Figure S2.** Stereomicroscopy images showing the *in vitro* dissolution of the different DMAP formulations after up to 3 min immersed in PBS.

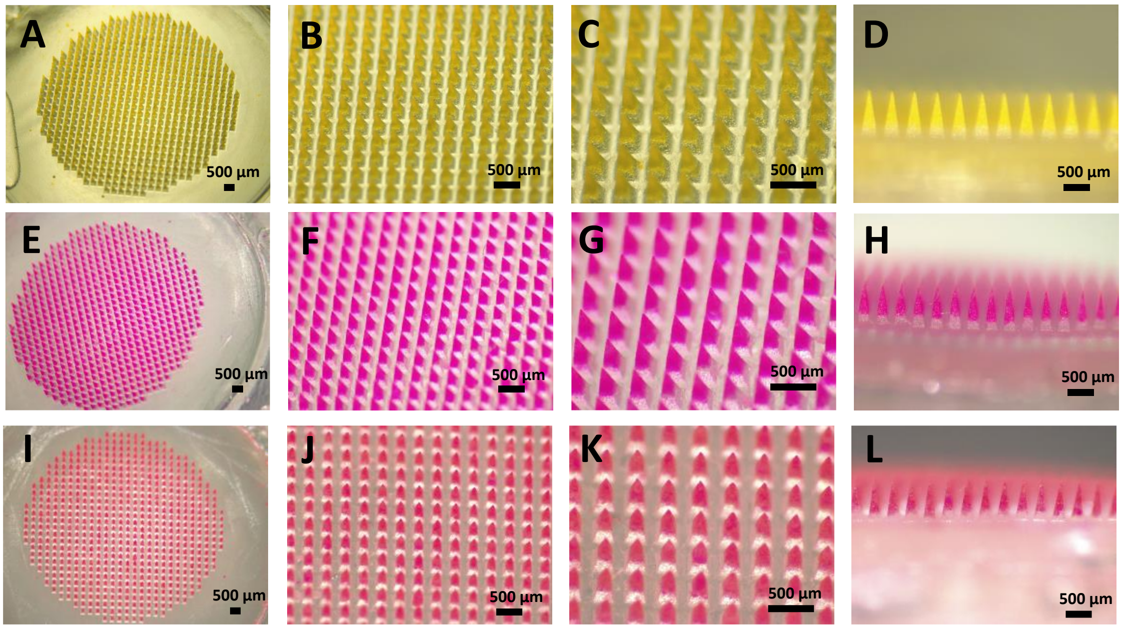


**Figure S3.** Stereomicroscopy images of DMAP containing FITC-labeled MSNs (A-D), RITC-labeled XL-MSN (E-H) or a combination of both types of MSNs (I-L).

**Table S1.** Characterization of the amount of fluorescently-labeled MSNs per DMAP.

| | Amount of FITC-M-MSN per MNA (mg) | Amount of RITC-XL-MSN per MNA (mg) | Total amount of MSN per MNA (mg) |
|---|---|---|---|
| FITC-M-MSN DMAP | 2.29 ± 0.06 | N/A | 2.29 ± 0.06 |
| RITC-XL-MSN DMAP | N/A | 2.31 ± 0.41 | 2.31 ± 0.41 |
| FITC-M-MSN + RITC-XL-MSN DMAP | 0.88 ± 0.05 | 1.56 ± 0.07 | 2.43 ± 0.08 |

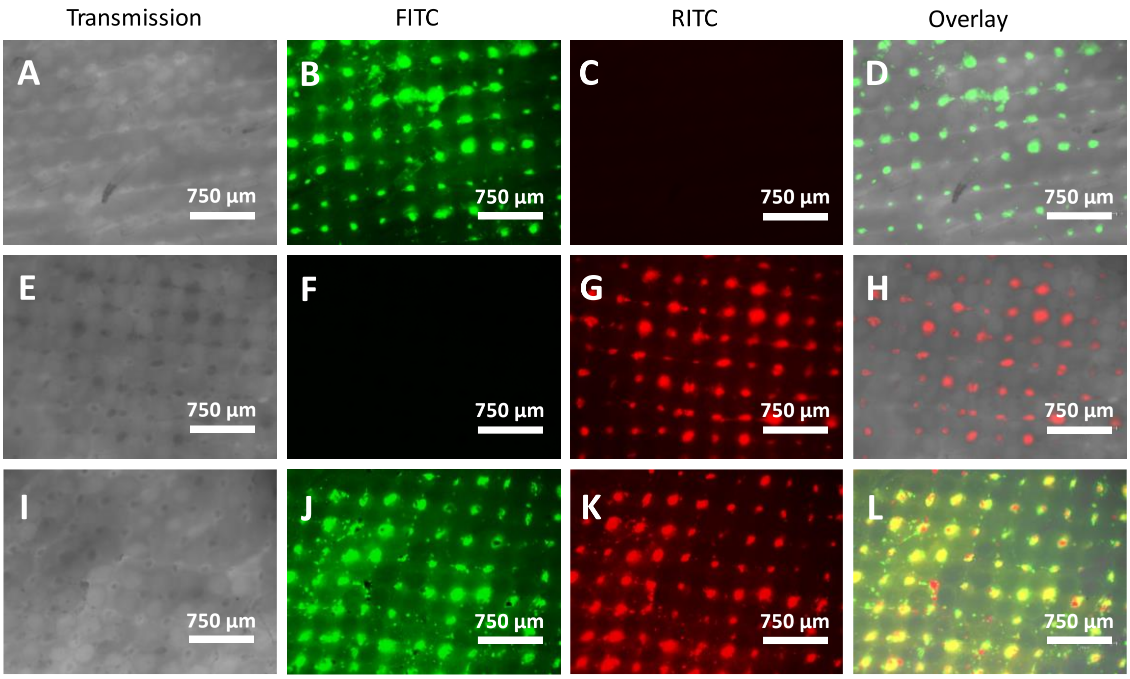


**Figure S4.** Fluorescence microscopy images of neonatal porcine skin after removal of DMAP containing FITC-labeled MSNs (A-D), RITC-labeled XL-MSN (E-H) or a combination of both types of MSNs (I-L).

**Table S2.** Evaluation of nanoparticle deposition of fluorescently-labeled MSNs per DMAP.

| | FITC-M-MSN Deposition % | RITC-XL-MSN Deposition % | Total MSN Deposition % |
|---|---|---|---|
| FITC-M-MSN DMAP | 20.88 ± 7.26 | N/A | 20.88 ± 7.26 |
| RITC-XL-MSN DMAP | N/A | 24.01 ± 14.02 | 24.01 ± 14.02 |
| FITC-M-MSN + RITC-XL-MSN DMAP | 21.184 ± 10.68 | 23.50 ± 9.23 | 22.72 ± 8.51 |

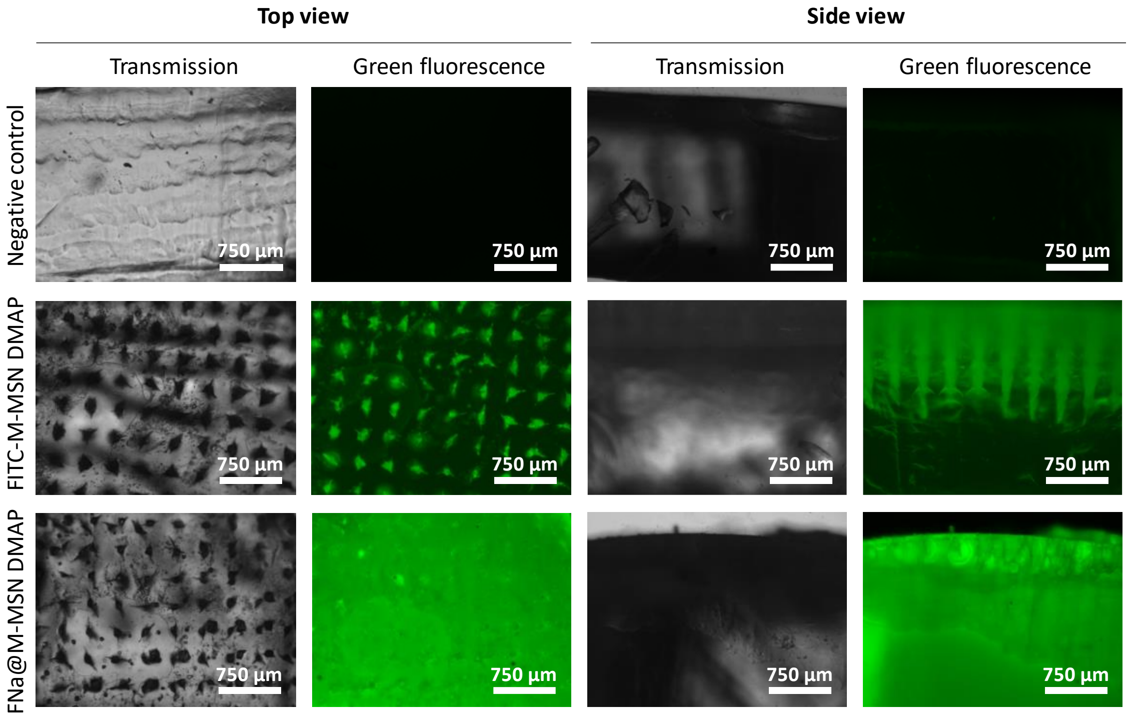


**Figure S5.** Fluorescence microscopy images of agarose gels 1 hour after insertion of DMAP without nanoparticles (top), containing FITC-labeled MSNs (center) and containing fluorescein sodium salt-loaded M-MSNs (bottom).

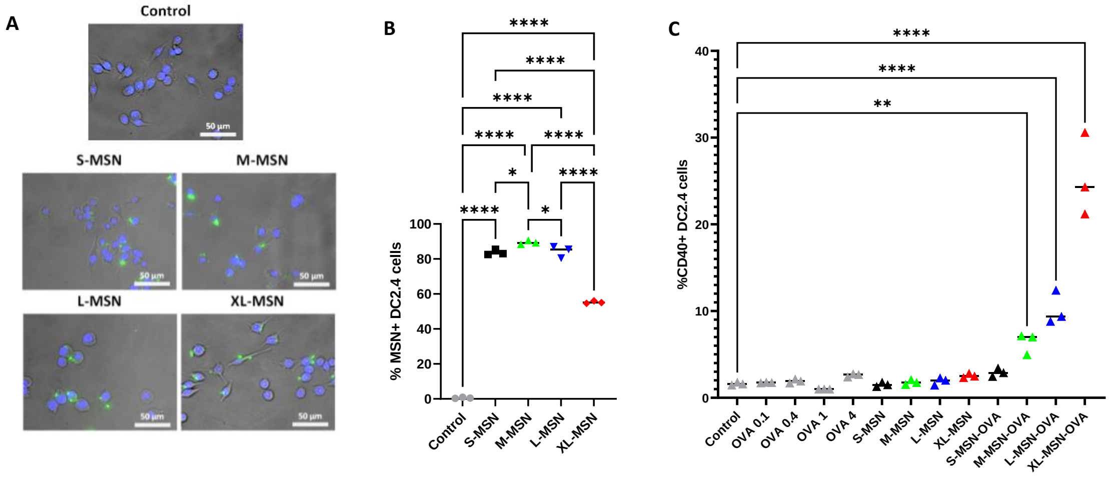


**Figure S6.** *In vitro* evaluation of MSNs incubated with DC2.4 cells. Fluorescence microscopy images of DC2.4 cells incubated with different types of FITC-labeled MSNs at a concentration of 10 µg/mL. Blue fluorescence from cell nuclei (DAPI), green fluorescence from MSNs (A); Flow cytometry results showing % of DC2.4 cells with nanoparticle uptake using different types of FITC-labeled MSNs at a concentration of 10 µg/mL (B); Activation of DC2.4 evaluated by expression of CD40 by flow cytometry upon treatment with free OVA (0.1-4 µg/mL), empty MSNs (10 µg/mL) and OVA-loaded MSNs (10 µg/mL). Data are Means ±SD, n=3. Statistical analysis by One way ANOVA. *p<0.05; **p<0.01; ***p<0.005; ****p<0.001.
